# A Landmark-based Common Coordinate Framework for Spatial Transcriptomics Data

**DOI:** 10.1101/2021.11.11.468178

**Authors:** Alma Andersson, Žaneta Andrusivová, Paulo Czarnewski, Xiaofei Li, Erik Sundström, Joakim Lundeberg

## Abstract

The increasing amount of spatial transcriptomics data prompts for means to amalgamate observations from distinct experiments, especially attractive is to cast quantities from different sources into a common coordinate framework (CCF) to relate signals across space. We here present a method that enables transfer of information from multiple samples to a reference representing a CCF, and show its utility by analyzing an assortment of real and synthetic data sets.

## 2 Main

During the last years, there’s been an ever increasing amount of interest in the field of spatially resolved transcriptomics, epitomized by its “Method of the Year 2020” award.[1, 2] The field has also experienced a trend of democratization, where techniques have spread beyond the groups originally developing them, a phenomenon reflected by the growing corpus of spatial transcriptomics data. Indeed, some of the spatial transcriptomics techniques have already been adopted as commercial products and embraced by the scientific community, thus facilitating the production of consistent high-quality data by a diverse set of labs. Spatial transcriptomics is also more frequently appearing as a modality of interest in ambitious international initiatives such as the Human Cell Atlas.[3] While quantity is key to delineate the many nuances of transcriptomics data, it also brings with it certain challenges; perhaps most notably the need to integrate observations from multiple sources.

For single cell transcriptomics data the concept of integration is often strongly associated with the process of constructing a shared space based on gene expression, to then embed the data therewithin. However, in contrast to single cell data, spatial transcriptomics data possess an inherent low-dimensional space, being the physical domain from which it’s collected. Thus, when building spatial transcriptomics atlases or summarizing larger studies, the idea of integration should be extended beyond elimination of unwanted batch effects. More specifically, it ought to encompass the transfer of data to a shared reference, where observations from different samples can be related in physical space. Such references are commonly referred to as common coordinate frameworks (CCFs), a concept which Rood et al. thoroughly discuss in their perspective.[4]

Considering this need for spatially aware integration methods, we here present a landmark-based approach to transfer spatial transcriptomics data to a defined reference. Our method relies on Gaussian Process (GP) regression, which previously has been successfully applied to identify spatially variable genes and cell interactions.[5, 6] With this method we seek to overcome both the limitations of traditional alignment methods relying on linear transformations (e.g., rotation and translation) as well as the need for an extensive preexisting reference system to which the data can be registered. We also provide an implementation of our method as a Python package, named “*effortless generic GP landmark transfer*”, or *eggplant* for short. To promote easy incorporation into already existing workflows and increase accessibility, *eggplant* is designed to be compatible with the popular analysis framework *scanpy* and its derivatives.[7]

To be more precise, our method focuses on the specific task of transferring observed spatial features from one coordinate system to a given reference system, using a set of shared spatial landmarks. The reference can be any arbitrary structure that represents a spatial domain onto which one seeks to transfer information, see Methods. Meanwhile, we define a spatial landmark as a feature that can be consistently located with fairly high precision across individuals. Samples where spatial landmarks (for brevity, we hereafter drop the prefix “spatial”) have been identified will be referred to as “charted”. Landmarks can be derived from any – to the tissue – associated information including morphological and molecular structures (e.g., gene expression or protein signals). Furthermore, the charting process can be manual, unsupervised (using computational methods) or a mixture of both; since our method is agnostic to this choice, we consider a deeper discussion regarding landmark annotation and identification to be outside the scope of this work. We also assume that the spatial data has been appropriately normalized and had eventual batch effects corrected for.

Our method is simple in its design and can be described in a few steps, see Figure 1A for a schematic overview. As input it requires charted spatial transcriptomics data containing one or more features of interest (FOI) together with a reference. The reference represents the domain to which the FOI’s distribution should be transferred and should also be charted. Next, the domain of the observed data is transformed to make landmark distances match those of the reference. The transformation can either be linear (multiplication with a scaling factor), or non-linear (using thin plate splines) if one suspects a non-homogeneous distortion of the spatial domain. Finally, we formulate a multivariate regression problem where the value of the FOI is considered a function of the distance to respective landmark. We employ a GP framework, commonly described as a distribution over functions, to learn the relationship between feature value and distances. A transfer of any FOI to the reference is seamless once the relationship is established; the function is simply applied to each location in the reference to obtain an estimate of the FOI value. Evidently, multiple samples can be transferred to the same reference, either one-by-one or jointly. Notably, there is no need for alignment or further processing once the samples have been charted. We also provide a strategy to determine a lower bound for the number of landmarks to be used in the process, see Methods.

**Figure 1:**
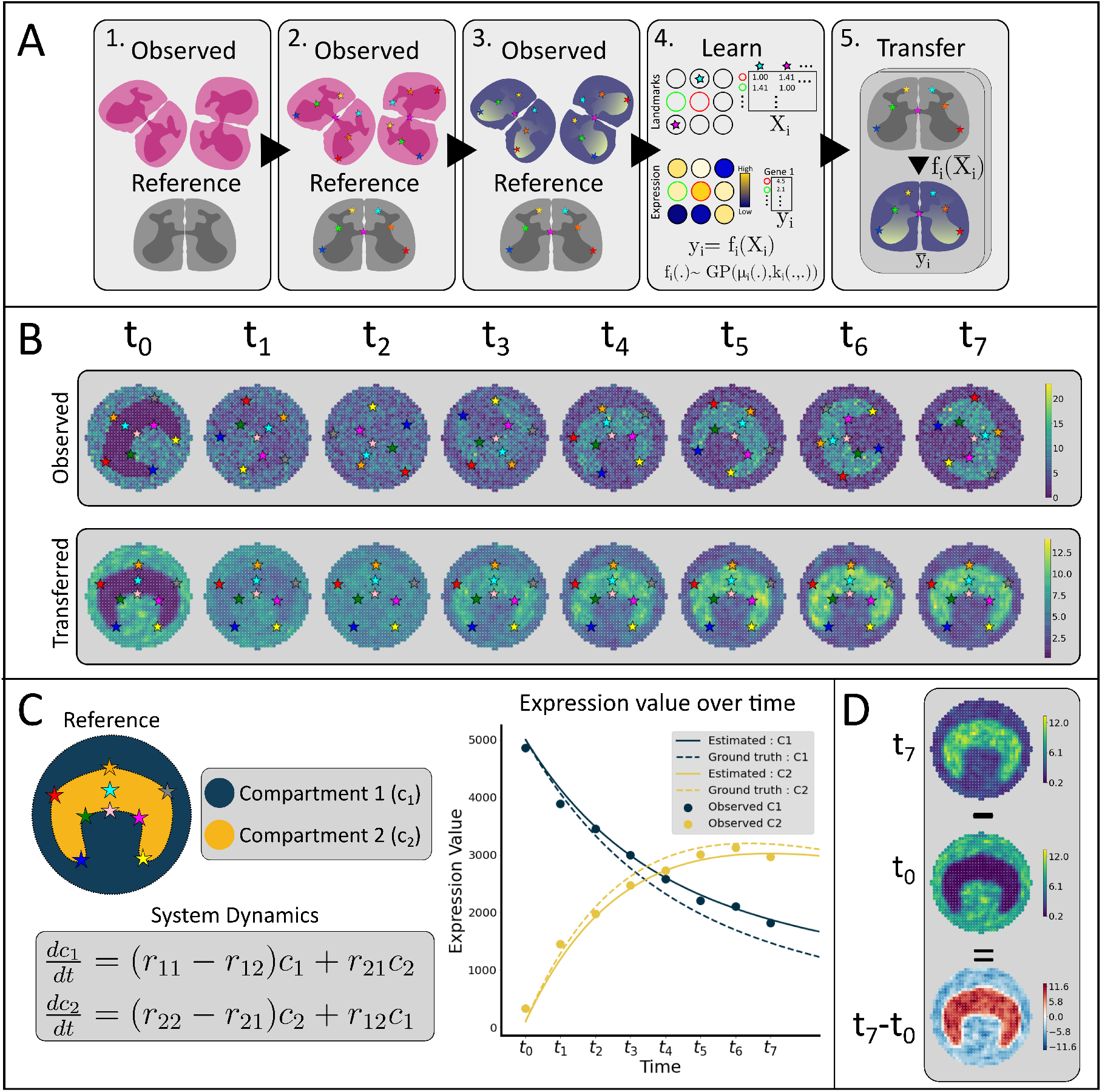
**A)** Schematic overview of the method. 1. We, select a number (here two) of samples representing the same spatial domain together with a reference. 2. We then chart the samples and reference (annotating landmarks). Here, landmarks are represented by colored markers. 3. Next, a feature of interest (FOI) that should be transferred to the reference is selected. 4. We learn the function that relates FOI values to landmark distances by using Gaussian Process (GP) Regression. 5. Finally, the FOI is transferred to reference using the learnt relationship between expression and landmark distances. **B)** Top : Observed synthetic data across eight different time points. Bottom : Results from transferring the observed data to a reference using our method. **C)** Spatiotemporal analysis of material (gene expression) transfer between the two compartments in the reference, the graph shows how the expression varies in each compartment as a function of time. **D)** An example of spatial arithmetics, subtraction of values at t_0_ from values at t_7_ shows local up-and downregulation of the feature between the two time points.

To demonstrate our method, we first apply it to a set of synthetic data containing eight samples from different time points in a dynamically changing system. The samples represent the same physical domain, but – like real data – exhibit differences in structure and orientation. Expression from each time point was transferred to a reference with the help of nine landmarks, Figure 1B. For this, and all subsequent analyses, we used non-linear landmark adjustment. This transfer of data to a CCF permits a multitude of downstream analyses, of which we will give two examples below.

The first example focuses on characterization of the system’s underlying spatiotemporal dynamics. The dynamical model used to generate the synthetic data is a two-compartment system, in which expression fluctuates according to a set of ordinary differential equations (ODEs). For the sake of simplicity, we assume that the model’s structure is known prior to the analysis, and therefore only aim to estimate the model’s parameters. The two compartments between which expression varies (C1 and C2) are defined in our reference, allowing us to approximate the total amount of expression in each compartment at every time point. From this aggregated data, we estimated the ODE-model parameters; the corresponding dynamics are shown in Figure 1C where they are also compared to the ground truth values. With the system dynamics established, we could also reconstruct the exchange of expression between the two compartments, see Supplementary Figure 1. In a biological system, this type of flux-analysis could for example elucidate how cells migrate between different regions in a tissue.

In a second example of downstream analysis, we leverage the fact that all data now inhabits the same reference, thus making coordinates comparable between time points. This allows us to perform “spatial arithmetics” from which information about local up-or downregulation of features between time points or conditions can be deduced and tested, see Figure 1D.

For additional evaluation of our method, a second set of synthetic data was generated to assess the influence of the number of landmarks on its performance and compare it to alternative strategies. In short, a non-homogeneous distortion was applied to a collection of spatial observations and associated landmarks, see Supplementary Figures 2A-B. We then assessed how well each strategy could recover the original spatial distribution of the distorted signals, where our approach exhibited the best performance, Supplementary Figure 2C. As expected, for landmark-based approaches, the number of landmarks was positively correlated with performance; however, this trend quickly diminished as the number of landmarks increased.

Having established confidence in our method, we next analyzed several sets of real spatial transcriptomics data. In the first analysis, we examined twelve first generation Spatial Transcriptomics (ST1K) samples of the mouse olfactory bulb (MOB), collected from different individuals and sexes.[8] Here we chose 14 landmarks, identified by morphological cues in the accompanying Hematoxylin and Eosin (HE) images, and charted the corresponding sites in our reference. Having prepared the data, we applied our method and transferred the expression of three genes to the reference: *Nrgn, Apoe* and *Omp*, see Figure 2A and Supplementary Figure 3 and 4. We also assembled “composite” expression profiles for each of the aforementioned genes, allowing us to represent information from all twelve samples jointly. We also conducted a “spatial differential expression analysis” (sDEA) between the three genes, to examine how their local expression differed. The composite representations and the sDEA results are both presented in Figure 2B.

**Figure 2:**
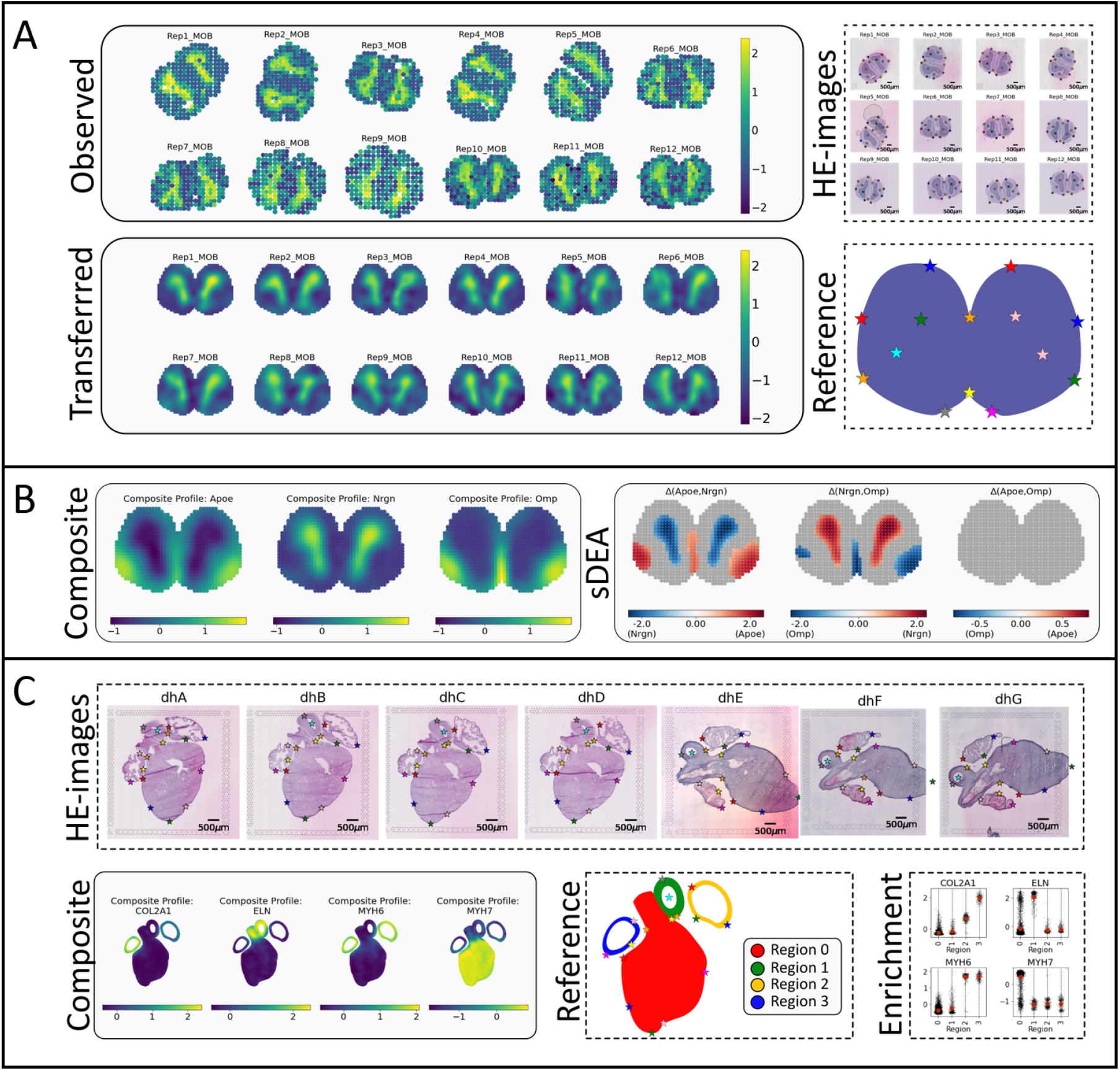
**A)** Top left: observed spatial gene expression of Nrgn in the mouse olfactory bulb (MOB) sections (n=12). Top right : charted HE-images of the MOB sections, landmarks are indicated by colored markers, for a larger image see Supplementary Figure 10. Bottom left: results from transferring the observed MOB data to a common reference. Bottom right : the charted reference to which the MOB expression data is transferred. **B)** Left : Composite profiles for each of the three genes Apoe, Nrgn and Omp. The composite expression profiles are formed by computing the location-wise mean across all twelve MOB sections, see Methods. Right : spatial differential expression analysis (sDEA) between the three genes, see Methods. Gray areas indicate locations where there’s no differential expression between the two compared genes. At locations with differential expression, the values for comparison Δ(g_1_, g_2_) are obtained by subtracting the composite profile of g_2_ from g_1_. **C)** Results related to the human developmental heart data. The “HE-images” panel shows the charted HE-images, landmarks are represented with colored markers. The ‘Composite” panel gives the composite representation (across samples, n = 7) of the transferred data for each gene. The “Reference” panel shows the reference to which data was transferred together with the four different regions, landmarks are indicated with colored markers. The “Enrichment” panel depicts the predicted expression values of each transferred sample (black dots) within respective region. Mean values are represented with a red marker.

In a second analysis, to show cross-platform compatibility, we also transfer gene expression in the mouse hippocampal area from data collected using both the Slide-seqV2 and Visium platforms. The Visium sample was charted with the help of the associated HE-image, while we relied on total UMI-counts for the Slide-seqV2 data, exemplifying how both morphology and molecular information may be used in the charting process (see Supplementary Figure 11). As shown in Supplementary Figures 13 (Observed) and 14 (Transferred), data from the two platforms were successfully integrated while preserving the intricate structure of the expression patterns.

Finally, we produced a new set of 10x Genomics Visium data consisting of seven sections (A-F) à two individuals from human developmental heart (dh) tissue (collected at the tenth postconceptional week). We then transferred the expression profiles of four genes (*COL2A1, ELN, MYH6* and *MYH7*) from all seven sections in this data set to a single reference. Despite vast inter-individual differences in the structure, the transferred data correlated well between patients; the mean between-individual correlation was 0.88, while the mean within-individual correlation was 0.96 for individual 1 and 0.91 for individual 2, see Supplementary Figure 15. We also generated gene-specific composite profiles, see Figure 2C. Separate representations of each combination of gene and section pairs are found in Supplementary Figure 5 (Observed) and 6 (Transferred). We also segmented the reference into four distinct spatial regions, which allowed us to assess region-specific enrichment of genes. Importantly, the enrichment analysis does not require any additional annotation of the original tissue samples, and the regions can be redefined without any need to repeat the transfer process. As expected, *MYH6* expression was highest in the atrial regions (Region 2 and 3), *MYH7* expression was elevated in the ventricular body (Region 0), and *ELN* in the outflow tract (Region 1).[9] The atria also were enriched for *COL2A1* but we, interestingly, observed a preserved and statistically significant left-right asymmetry in its expression (*p*_*value*_ *<* 0.05, two-sided permutation test).

Gene expression may be the primary information that spatial transcriptomics techniques produce, but there’s now a panoply of methods to infer second order insights from said data. Thus, to demonstrate the flexibility of our method, we transferred inferred (by the tool *stereoscope*) cell type proportion values between two Visium sections of human breast cancer, see Supplementary Figure 7.[10]

In this study we have presented a new and general method to transfer spatial transcriptomics data from multiple samples to a shared reference, something that previously only has been conceptually described. The method is versatile and effortless to use. Furthermore, the implementation leverages the GPyTorch framework, which supports GPU acceleration together with efficient algorithms to reduce the inference’s complexity.[11] Our tool is useful for visualization purposes, but also prepares the data for more extensive analysis, such as spatiotemporal modeling, spatial arithmetics, and regional enrichment. We are currently relying on manual identification of landmarks, but see a great opportunity for future research to explore different venues for unsupervised landmark detection. Taken together, we consider this an important first step towards harmonizing and integrating spatial transcriptomics data in a common coordinate framework.

## 3 Methods

### 3.1 Code Availability

An implementation of our method is provided as a Python package named *eggplant*, short for ***e****ffortless* ***g****eneric* ***GP lan****dmark* ***t****ransfer*. The package can be accessed at the GitHub repository https://github.com/almaan/eggplant. The repository also contains a set of Jupyter notebooks outlining all the presented analyses as well as generation of the synthetic data associated with this study. The repository also contains scripts to download and curate the public data that we’ve used. We have also deposited a clone of the repository together with the charted data at Zenodo, accessible via https://doi.org/10.5281/zenodo.5659093.

### 3.2 Data Availability

Except for the synthetic and developmental heart data, we used publicly available data sets in this study. We thus refer to the original data sources for access, which we list below:

- Synthetic data: https://github.com/almaan/eggplant
- MOB data: https://www.spatialresearch.org/resources-published-datasets/doi-10-1126science-aaf2403/
- Hippocampal region Visium: https://support.10xgenomics.com/spatial-gene-expression/datasets/1.1.0/V1_Adult_Mouse_Brain
- Hippocampal region Slide-seqV2 (Puck_200115_08): https://singlecell.broadinstitute.org/single_cell/study/SCP815/highly-sensitive-spatial-transcriptomics-at-near-cellular-resolution-with-slide-seqv2
- bcA: https://support.10xgenomics.com/spatial-gene-expression/datasets/1.1.0/V1_Breast_Cancer_Block_A_Section_1
- bcB: https://support.10xgenomics.com/spatial-gene-expression/datasets/1.1.0/V1_Breast_Cancer_Block_A_Section_2
- Single cell HER2 data : https://zenodo.org/record/4739739#.YPF2D5KxVhE

For the human developmental heart, raw sequencing data can be accessed at the Gene Expression Omnibus (GEO) with access code GSEXXXXXX (*). All processed data together with the presented results are available at the GitHub and Zenodo repositories associated with this manuscript.(*)

*\*Note*: all new data will be publicly available upon publication of the manuscript.

### 3.3 Data Acquisition

#### 3.3.1 Human Developmental Heart

After collection, the human developmental heart tissue was fresh-frozen and embedded in Tissue-Tek (OCT). The tissues samples were cryosectioned at 10 *µ*m thickness and placed on 10X Visium spatial gene expression slides, to then be stored at − 80^*°*^*C* prior the library preparation. Libraries were generated from the samples using Visium Spatial Gene Expression kit from 10x Genomics. Every barcoded Visium array contains four capture areas á 4992 spots, where each spot contains probes consisting of: a spatial barcode, an UMI sequence, and a poly-dT-VN sequence enabling mRNA capture. Sections were fixed for 30 min in Methanol, stained with Hematoxylin and Eosin and imaged using Metafer Slide Scanning system (Metasystem, Altlussheim, Germany). The 10x Genomics Visium Tissue Optimization Kit was used to determine the optimal permeablization time, rendering an estimate of 20 mins. The generated libraries were sequenced using the Illumina Platform. The lengths for read 1 and read 2 were 28 bp respectively 120 bp. The sequencing data was processed with *spaceranger* v.1.2.0.

### 3.4 Data Processing

In the Slide-seqV2 data, we removed all beads with less than 100 UMI’s and then subsampled the remainder to 20% of its size. For Visium and first generation Spatial Transcriptomics (ST1K) data, we used all spots identified to be under the tissue (for public data sets we used the original annotations).

When analyzing gene expression data, we applied a simple normalization strategy compiled of functions from the *scanpy* (v. 1.8.1) package, the recipe is given below:

1. *scanpy*.*pp*.*filter_genes(*…,*min_cells = 0*.*1)*
2. *scanpy*.*pp*.*normalize_total(*…,*1e4, exclude_highly_expressed = True)*
3. *scanpy*.*pp*.*log1p(*…)
4. *scanpy*.*pp*.*scale(*…)

When cell type proportions acted as the feature of interest we only used standard scaling (subtraction by mean and division by standard deviation).

Working with the older ST1K data, we also added a *spatial smoothing* step to the above recipe (as a last step), to counteract “holes” caused by tears or ruptures of the tissue as well as steep gradients and variation in the capture efficacy across the tissue. The spatial smoothing is a form of weighted average of the feature values observed in a given location’s neighborhood. The neighborhood of spot *s* is denoted as 𝒩 (*s*) and contains said spot together with its four nearest neighbors. If *y*_*s*_ is the prior feature value associated with spot *s*, then the smoothed equivalent 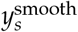 is defined as:

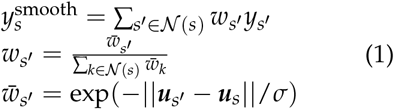

Where ***u***_*s*_ are the coordinates of spot *s* and ||.|| represents the L2-norm (euclidean distance). In our analysis we used *σ* = 50.

### 3.5 Model

The method we propose transfers a feature of interest from one coordinate system to a given reference system, below we describe the process in more detail. Let Ω be the domain from which the observed data is collected, while Ω′ represents the reference domain onto which we seek to transfer information. Similarly, ℒ ⊂Ω is the set of landmarks in the observed data, while ℒ ′ ⊂Ω′ represents the landmark positions in the reference. Here, |ℒ| = |ℒ ′| = *L*, where *L* is the number of landmarks and |.| is the cardinality operator. Importantly, ℒ and ℒ ′ are ordered in the same way. We also define 𝒰 ⊂Ω and 𝒰′ ⊂Ω′ as the sets of coordinate tuples containing the location of each observation (***u***_*i*_) and reference points 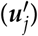. Every observation *i* has a target value *y*_*i*_ associated with it, and our primary objective is to find the corresponding values 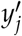 for the reference points.

First, we will transform the coordinate tuples in 𝒰 and ℒ, to put distances between objects in the two sets at the same lengthscale as between their reference counterparts (𝒰′ and ℒ ′). The transformation *h* can either be a simple linear scaling: *h*(***u***_*i*_) = *h*_const_(***u***_*i*_) = *a ·* ***u***_*i*_, or a more complex transformation relying on thin plate splines (TPS). In the case of the former, *a* will be given as the average of ratios between landmark-pair distances, that is:

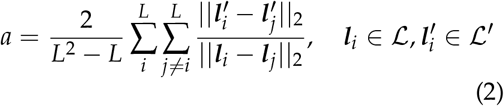

In the second case, *h* will be a composite function given as *h*(***u***_*i*_) = *h*_TPS_(*h*_const_(***u***_*i*_)), where *h*_TPS_ restricted to a family of TPSs parametrized by minimizing the cost *C*:

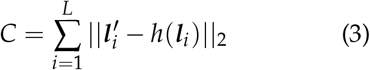

The transformed versions of 𝒰 and ℒ, obtained by applying *h* to every element in respective set, are referred to as 𝒰^*^ and ℒ^*^. In our implementation, we use the Python package *Morphops* (v. 0.1.12) for the TPS warping.

Next, for all members of 𝒰^*^ and *U′*, we compute the distances to ℒ^*^ respectively ℒ ′, forming the two new sets *X* and *X′* defined as:

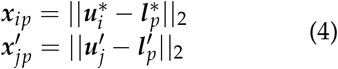

We then seek a function *φ* that will allow us find the feature values associated with each location in our reference:

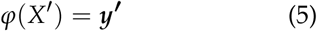

To learn *φ*, we use Gaussian Process Regression (see Section 3.6) where the observed data is used to learn said function.

### 3.6 Gaussian Process Regression

Gaussian Process (GP) Regression is fundamental to our method, and we will therefore briefly describe it in the context of our work. However, for a more elaborate account of GP regression we refer to any of the (many) already existing works on the subject, for example the canonical text by Rasmussen and Williams.[12]

A GP is defined as a collection of random variables, of which any finite subset have a joint Gaussian distribution. Hence, a GP may be interpreted as a distribution over functions that fit a certain set of points. We denote a function *f* that is distributed according to a GP as:

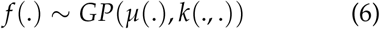

Here, *µ*(.) and *k*(., .) represent the mean respectively covariance function (also referred to as the kernel).

In our model, the function f relates landmark distances to the feature of interest’s values. We represent the complete set of observed data In our model, the function *f* relates landmark distances to the feature of interest’s values. We represent the complete set of observed data as the tuple (*X*, ***y***), where *X* ∈ ℝ^*M*×*L*^ is the matrix representing the distances to each of the *L* landmarks for all of the *M* observations, and ***y ∈*** ℝ^*M*^ is the value of the feature of interest associated with each observation. Distances and feature values are related via *f*, that is *f* (*X*) = ***y***. The distances from the locations to the landmarks (in the reference) are represented by *X′* while ***y′*** indicates the reference target values (which we seek to approximate).

Due to the properties of GPs, the joint distribution *p*(***y, y****′* |*X, X′* ; *σ*) thus becomes:

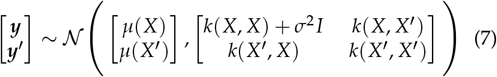

Where we account for noise in the training data according to the model : ***y*** = *f* (*X*) + *ε, ε* ∼ 𝒩 (0, *σ*^2^). Using standard Gaussian identities and the assumption *µ*(.) = *c ·* **1** = ***c***, where *c* is some real number, the conditional distribution *p*(***y′*** |*X′, X, y*; Θ) becomes:

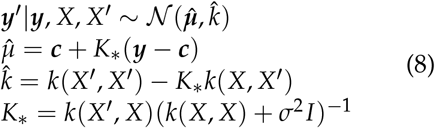

With (*X*, ***y***) being given, we consider the conditional mean a function of *X′* and will use this as *φ* described in Section 3.5, that is:

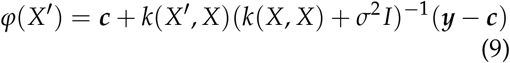

We support several different kernel functions but use the RQKernel as default, which is defined as:

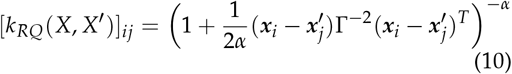

Where ***x***_*i*_ and 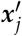 refers to the i:th respectively j:th row of *X* and *X′*, while *α ∈* ℝ and Γ ∈ ℝ^*L*^ are kernel parameters. To find optimal values of the parameters Θ = [*c, σ, α*, Γ], we optimize the marginal likelihood *p*(***y*** *X*; Θ) using stochastic optimization. Once these parameters have been estimated, *φ* can be used to estimate ***y****′*. Implementation-wise we leverage the GPyTorch (v. 1.5.0) framework for both inference and prediction.

### 3.7 Synthetic Data

Here we outline the process by which each synthetic data set was created, the time series data refers to the set analyzed in Figure 2 while the distortion data refers to the set presented in Supplementary Figure 2.

#### 3.7.1 Time series data

Eight two-colored images (see Supplementary Figure 9) were used to generate the spatial domain for each time point. To convert images to array data, *eggplant*’s *reference_to_grid* function from the *preprocess* module was used, this also assigned each spot in the array to one of two groups (Compartment 1/C1 and Compartment 2/C2). The number of transcripts in each compartment was dictated by the dynamical system; from which expression values at select time points were extracted and rounded to the nearest integer value. The transcripts were then randomly distributed between array nodes in the associated compartment. Below we describe the dynamical model in more detail.

The dynamical model describes a two-compartment system governed by the following set of equations:

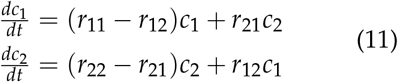

Where *c*_1_ is the amount of material in compartment 1 and *c*_2_ the same but for compartment 2. From a given set of initial values, (*c*_1_(0), *c*_2_(0)), the system was then propagated in time for a pre-determined number of steps (*T*). Here the following parameter values – arbitrarily chosen – were used: (*r*_11_, *r*_12_, *r*_21_, *r*_22_) = (0.2, 0.1, 0.8, 0.3) together with the initial values (*c*_1_(0), *c*_2_(0)) = (5000, 100). The eight time points from which we extracted expression values were equally spaced in in the interval [0, 500].

In figures, tables and text we refer to this synthetic data set as “Synthetic 1”.

#### 3.7.2 Distorted data

First, a *p* × *p* grid where each node represented a spatial capture location (e.g., spot) was generated, to figure as the domain in which signals will be collected. Next, to produce a spatial expression pattern, an *i* iterations long random walk was performed (the initial position also being randomly sampled from the domain). The number of times a node was visited in the walk was let to represent its observed expression level; this data represent the “ground truth”. From the ground truth, a “distorted” representation of the same sample was produced by first applying a distortion field (*F*(*x, y*)) to the node positions while keeping their values constant. Then, we placed a new *p* × *p* grid identical to the first over the distorted data, and interpolated its node values by a nearest neighbor approach. For a depiction of the process see Supplementary Figure 2. For our data we let *p* = 32, *i* = 1 × 10^4^, and 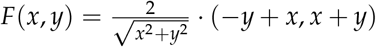

In figures, tables and text we refer to this synthetic data set as “Synthetic 2”.

### 3.8 Choosing the number of landmarks

While including more landmarks generally will render a better result, this gain in performance tends to be marginal after a certain number of landmarks have been included in the analysis. Hence, we aim to provide means to estimate a *lower bound* of the number of landmarks that should be used when transferring information to a reference. Below, we describe the steps to derive this lower bound.

First, we select one representative sample from our data set and position *L* landmarks in the (spatial) domain which the sample inhabits. Then, landmarks are randomly placed in the domain using Poisson Disk Sampling, where the first landmark always is located at the domain center.[13] We denote the set of all landmarks as ℒ, this set is considered as ordered. Next we specify a sequence (*N*_*L*_ = {*l*_1_, ..*l*_*P*_}, *l*_1_ ≥1, *l*_*p*_ ≤*L*) of the numbers of landmarks that should be evaluated. Then, for each entry *l*_*i*_ we randomly choose *l*_*i*_ of the *L* landmarks, and learn the transfer function using the representative sample. In this analysis, the normalized total-UMI count figures as the feature of interest. For each number of landmarks *l*_*i*_, we compute the mean negative marginal log likelihood (nMLL) for the last *T* iterations when fitting the model, and compare the mean values between all numbers in *N*_*L*_. This process is repeated for *n*_*rep*_ times, which allows us to compute an average for each element *l*_*i*_. By inspecting the graph obtained by plotting the average nMLL values as a function of the number of landmarks and applying a *Savitzky–Golay* filter for smoothing, we let the lower bound be defined as the number of landmarks where the average nMLL starts to plateau.

Table 1 shows the estimated lower bounds together with the actual number of used landmarks in the analysis, the graphs from which the lower bounds were determined are shown in Supplementary Figure 16. In all of our analyses we aimed to use as many landmarks as we could confidently identify, with the requirement that this number should be higher than the – to each sample – associated lower bound.

**Table 1:**
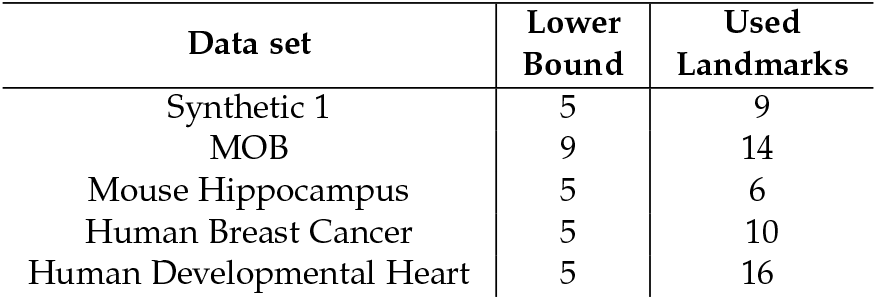
Landmark lower bounds and number of used landmarks. The column “Lower bound” gives the estimated lower bound for the number of landmarks to be used in each data set. The column “Used Landmarks” lists the number of landmarks actually used in the analysis. The representative sample (S) from each data set (D) are given as (D,S): (Synthetic 1, t_7_), (MOB, Rep1), (Mouse Hippocampus, Visium), (Human Breast Cancer, bcA), (Human Developmental Heart, dhA).

| Data set | Lower Bound | Used Landmarks |
| --- | --- | --- |
| Synthetic 1 | 5 | 9 |
| MOB | 9 | 14 |
| Mouse Hippocampus | 5 | 6 |
| Human Breast Cancer | 5 | 10 |
| Human Developmental Heart | 5 | 16 |

The following parameter values were used: *N*_*L*_ = {1, 3, 5, 7, 9, 11, 13, 15, 17, 20}, *T* = 200, *n*_*rep*_ = 5, for the Savitzky–Golay filter we used the function *savgol_filter* from the *scipy*.*signal* module (v. 1.7.1) with parameters `window_length=5` and `polyorder=4`.

### 3.9 ODE parameter estimation

To estimate the parameters of the ODE system representing the dynamical model, after the synthetic data had been transferred to the reference, we used the BFGS algorithm with a cost function dependent on the system model (Equation 11). First we aggregated the data in each compartment to get an expression tuple for every time point, that is:

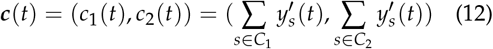

Where *C*_*i*_ is the set of spots in compartment 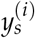 is the transferred expression value at array point *s*, and *t* represents time point *t*. Next, let *p*(., ***r***; *T*) represent a function that propagates the first argument according to the dynamics given in Equation 11 *T* steps forward in time with parameter values ***r***. From this, the cost (*C*) for a given set of parameters ***r*** takes the form:

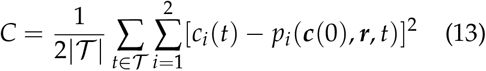

Where 𝒯 is the set of observed time points. We used the *minimize* function from *scipy*’s *optimization* module for the optimization, and *odeint* function from the *integrate* module to solve the ODE system; with *scipy* (v. 1.7.1)

### 3.10 Spatial Arithmetics

Conducting any form of spatial arithmetics is straightforward once observed data has been transferred to the same reference. If we let *λ*(., .) represent an arbitrary arithmetic operation, and 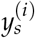 denotes transferred data from sample *i* at location *s* in the reference, then:

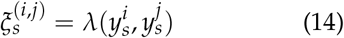

gives the expression for the spatial arithmetic calculation, where 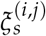 is the output associated with location *s*.

### 3.11 Spatial Differential Expression Analysis

From the Gaussian Process Regression, we obtain both mean and variance estimates of the feature values at each location, together these can be used to perform *spatial differential expression analysis* (sDEA) between groups of samples (e.g., disease vs. control). First, we compute the local group mean (*µ*) and variance (*σ*^2^) values for a FOI, which for any group *G* and location *s* are defined as:

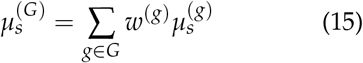

And

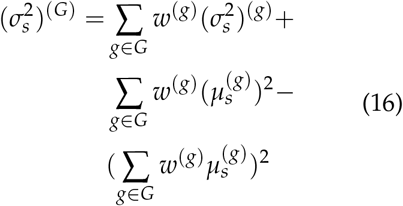

Where *w*^(*g*)^ denotes the weight that should be given to sample *g* when computing the mean, and 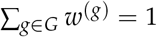 if nothing else is stated we assign equal weights to all samples within the same group. Next, for each group *G* and location *s* we construct an interval 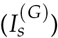 according to:

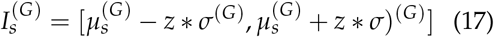

Where *z* relates to the number of samples that would fall into the interval if we were to sample new values from the mixed distribution, if nothing else is stated we use *z* = 2. Finally, we consider the FOI to be to be spatially differentially expressed at location *s* between the two groups *G*_*i*_ and *G*_*j*_ if the two intervals 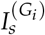 and 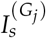 do not overlap. Evidently, a larger value of *z* will require that the two groups are more distinct in their expression of the FOI to be considered spatially differentially expressed at a given location.

### 3.12 Analysis

#### 3.12.1 Transfer to reference with *eggplant*

In all analysis steps we used 1000 epochs, an *RQKernel*, and the Adam optimizer with a learning rate of 0.01. The references were all represented by approximately 1000 array points, except for the human developmental heart data where 10000 array points were used. The number of landmarks used in each analysis are listed in 1. The landmarks did not correspond to any “established” anatomical features but were rather selected based on their ease of identification from the morphology or gene expression pattern across the examined samples.

All references used in our analyses are found in Supplementary Figure 8 together with their respective landmark annotation. The charted observed data is displayed in Supplementary Figure 9-12. All this information is also available in the – to this manuscript –associated GitHub repository.

#### 3.12.2 Benchmarking and Landmark Influence

We compared the transfer made by *eggplant* with three alternative strategies: “no correction”, “constant mean”, and thin plate spline interpolation (TPS). The task designed to measure performance consisted of trying to transfer distorted data back to its original (ground truth) distribution in a data set generated according to the procedure described in Methods Section 3.7.2. The Root Mean Squared Value (RMSE) value between the ground truth and the corrected values was used as a metric to assess performance. In the “no correction” strategy, the grid values in the distorted data is immediately compared to the ground truth values. This strategy emulates a scenario where tissue sections would be aligned, but non-linear distortions not accounted for. In the “constant mean” approach, we assign all grid points the same value, being the mean value. Notably, the expected RMSE value for this approach is 1 since we applied standard scaling to the data. Finally, with the TPS method, the same landmarks as provided to *eggplant* were used to correct for the distortion; then every grid point in the reference domain was assigned the value of its nearest neighbor among the shifted data points. We compared the strategies with different number of landmarks (*L* ∈ {3, 7, 11, 15, 19, 23, 27, 31, 35, 39}) and repeated each comparison 3 times to asses variance of the outcome. In each iteration, the landmarks were selected from a set of 40 random positions – sampled by the same Poisson Disc Sampling strategy as referenced above – in the spatial domain and then distorted by the same field *F* as the grid points during generation of the distorted set. For the TPS strategy we use the *Morphops* (v. 0.1.12) package, for the 2d interpolation we used *scipy*.*interpolate*’s *griddata* function (v. 1.7.1).

#### 3.12.3 Statistical Tests

In our study we perform a permutation test to asses whether there’s an asymmetry between the two different atria (Region 2 and 3) w.r.t. *COL2A1* expression in the human developmental heart data set. We favored a permutation test since our observations violate the i.i.d. assumption that most statistical tests rely on. We outline how this test is constructed below.

For two arbitrary regions A and B, we let *R*_*A*_ and *R*_*B*_ denote the sets of feature values associated with the locations contained within respective region. Without loss of generality, we here assume that our objective is to determine whether the expression of a feature of interest differs between region A and region B. We define the *mean region difference* (Δ_*A,B*_) as the mean of the difference in feature value across all combinations of observations from each set. That is:

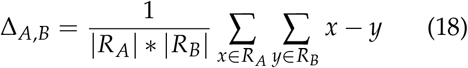

Our objective is then equivalent to testing whether the observed mean region difference is more extreme than what is expected by chance. To perform this test we shuffle the observations’ region labels and compute the Δ_*A,B*_ value for each permutation. We then compute the p-value as:

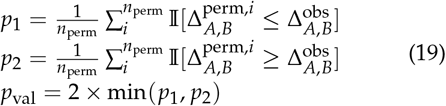

Where 𝕀 is the indicator function. If *p*_val_ ≤ *α*, the difference in expression between the two regions is considered statistically significant. Here *α* is the significance level, and the test is two-sided in its character. In our analysis of the *COL2A1* right-left asymmetry, we let *α* = 0.05 and ran 1000 permutations.[14] The test was applied to the *composite* representation of the *COL2A1* expression.

#### 3.12.4 Single cell mapping with *stereoscope*

For the *stereoscope* (v. 0.3.1) analysis we used the *major* cell type tier found in the single cell data, only including cells from HER2-positive patients. Cell types with less than 25 members were excluded, for cell types with more than 500 members, a subset consisting of 500 cells were randomly sampled from these. We also used a curated list of genes in the analysis consisting of 5540 members, representing a union of the 5000 highest expressed genes and cell type specific marker genes, see Supplementary Data 13 in [15]. We used 50000 epochs and a batch size of 2048 for the single cell parameter estimation as well as the proportion inference.

## 3.13 Contributions

A.A. conceived the method, implemented it in code, analyzed the data, generated the results, and wrote the paper under supervision of J.L. P.C. was involved in the discussion of using a landmark-based approach, and also beta-tested the code. Together, Ž.A., X.L., and E.S. produced the human developmental heart data, Ž.A. also helped in the writing of the experimental methods part. All authors read and gave feedback on the paper.

## 3.14 Acknowledgments

The authors thank Marco Vicari for important feedback and discussions regarding the developmental heart’s anatomy and structure. We also thank both Franziska Hildebrandt and Ludvig Larsson whom read the paper and provided constructive feedback.Finally, a special tanks is directed to Ludvig Bergenstråhle who gave invaluable feedback. This work was made possible by generous support from the Knut and Alice Wallenberg foundation, the Erling-Persson family foundation, the Swedish Cancer Society, the Swedish Foundation for Strategic Research, Karolinska Intsitutet Research Funds, and the Swedish Research Council. Human developmental heart tissue was acquired through the Karolinska Institutet Developmental Tissue Bank.

## 3.15 Ethics declaration

The use of human developmental heart in the study was approved by the Regional Ethical Review Board in Stockholm and the National Board of Health and Welfare. The procuration of the tissue and processing of the data were in concordance with the ethical stipulations of the Helsinki Convention (Dnr: 2:9/2015). The human developmental heart tissue used in this study was retrieved from medical abortions at the Department of Gynecology, Danderyd Hospital and Karolinska Huddinge Hospital, all patients provided a written statement of informed consent.

