## supplementary material for "A Landmark-based Common Coordinate Framework for Spatial Transcriptomics Data"

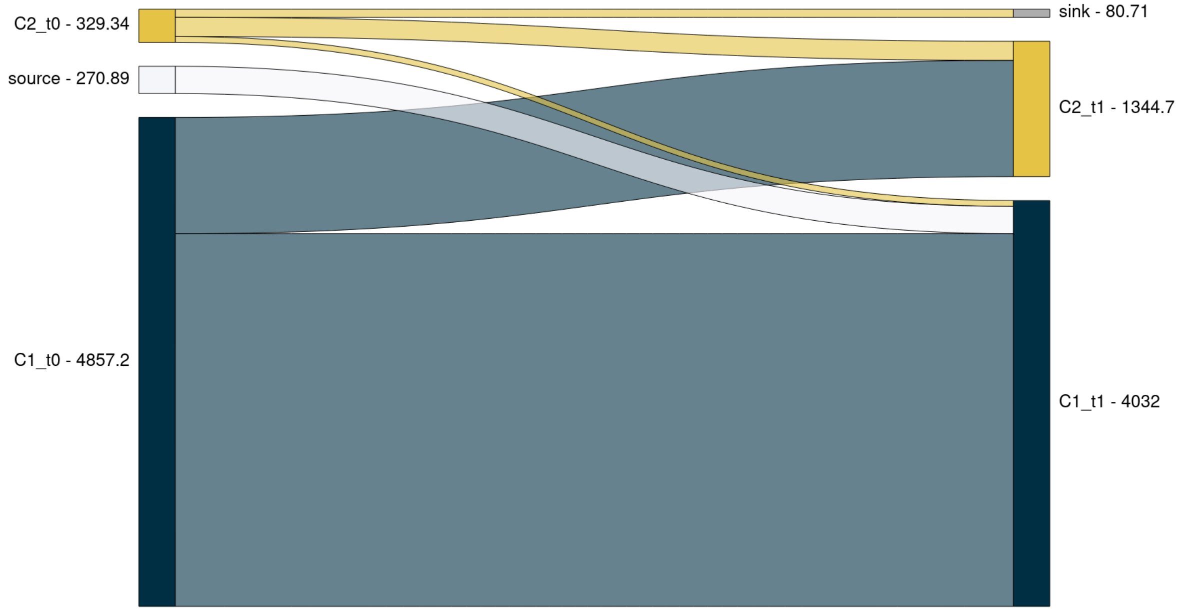


**Supplementary Figure 1:** *Sankey Diagram representing the net flow of material from each compartment. The label "source" represents addition of new material to the system (exclusive to C1), while sink represents loss of material from the system (exclusive to C2). The numbers represent the total amount of material in each compartment at respective time point. Labels are given as CX_tY, X = Compartment and Y = time point.*

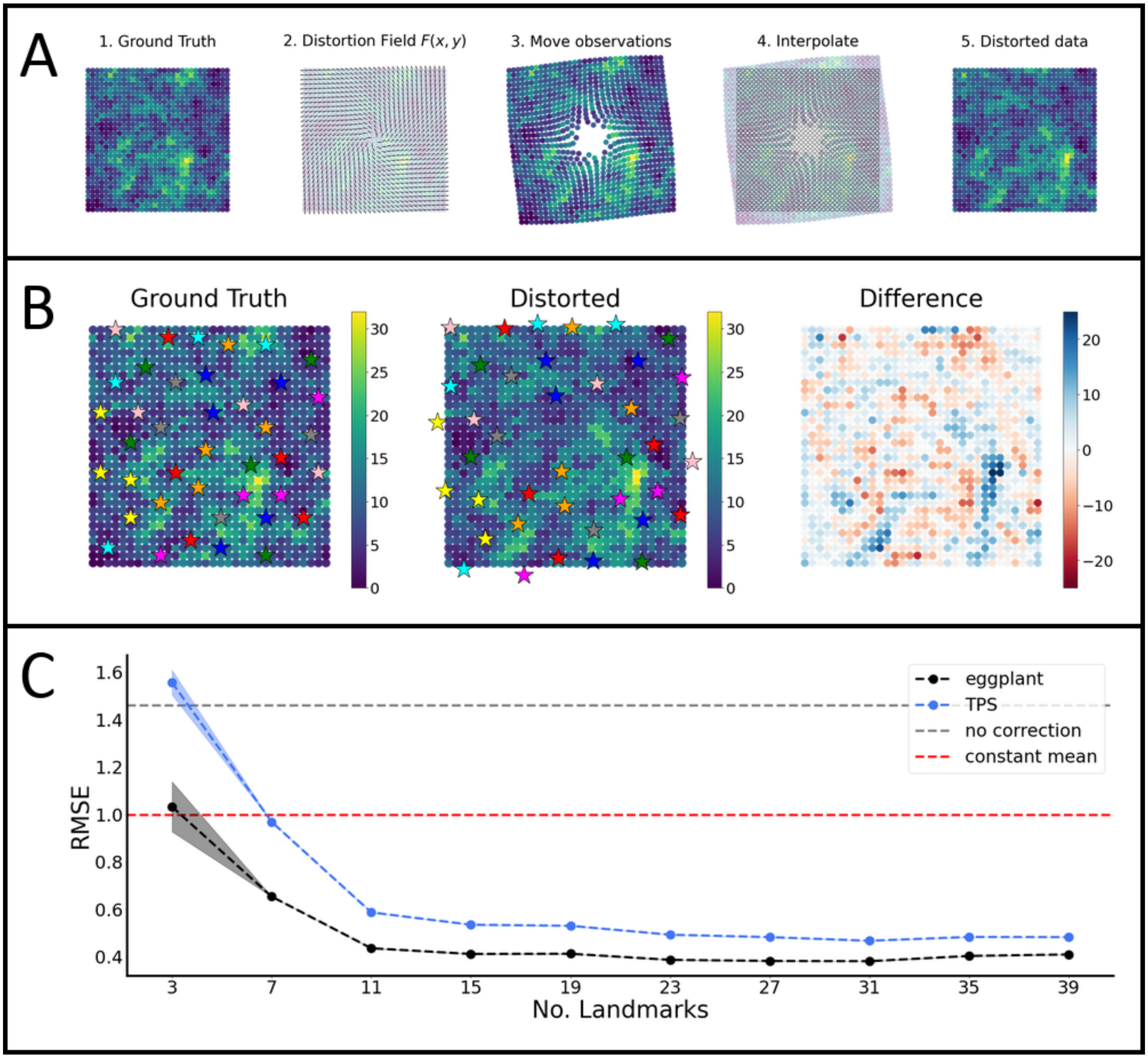


**Supplementary Figure 2:** ***A)*** *Overview of the process designed to generate distorted data from the ground truth data. First (1), ground truth data is generated by a random walk strategy. Next (2-3), a distortion field is applied to the nodes in the grid, perturbing the grid points' spatial location. Then (4), a grid identical (in structure) to the ground truth's is overlaid on the shifted data, (5) to which the values from the distorted data are assigned based on their nearest neighbor in physical space.* ***B)*** *Overview of the ground truth and distorted data together with the* 50 *landmarks from which a subset were taken in each analysis. Here, the "Difference" subplot represents the difference in signal strength between the locations in the ground truth and distorted data, (former subtracted from the latter).* ***C)*** *Influence of the number of landmarks on the performance of four different strategies for estimation of the original spatial distribution (ground truth) from the distorted data. The shaded regions indicate* $\pm 1$ *standard error (SE). TPS is short for "Thin Plate Splines"*

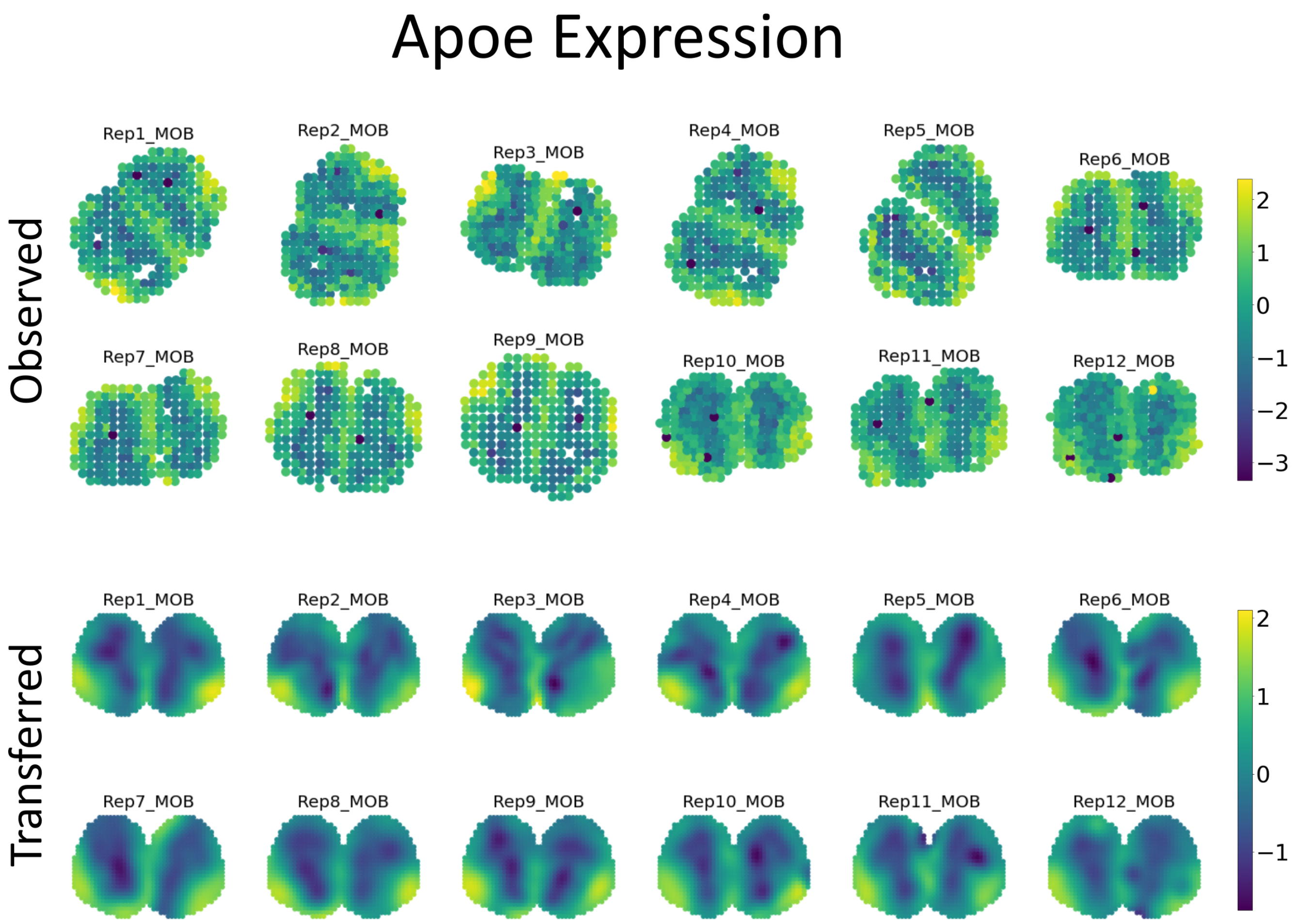


**Supplementary Figure 3:** *Top : Normalized observed spatial gene expression of Apoe in the twelve mouse olfactory bolb (MOB) sections. Bottom : results from transferring the observed spatial gene expression to a shared reference.*

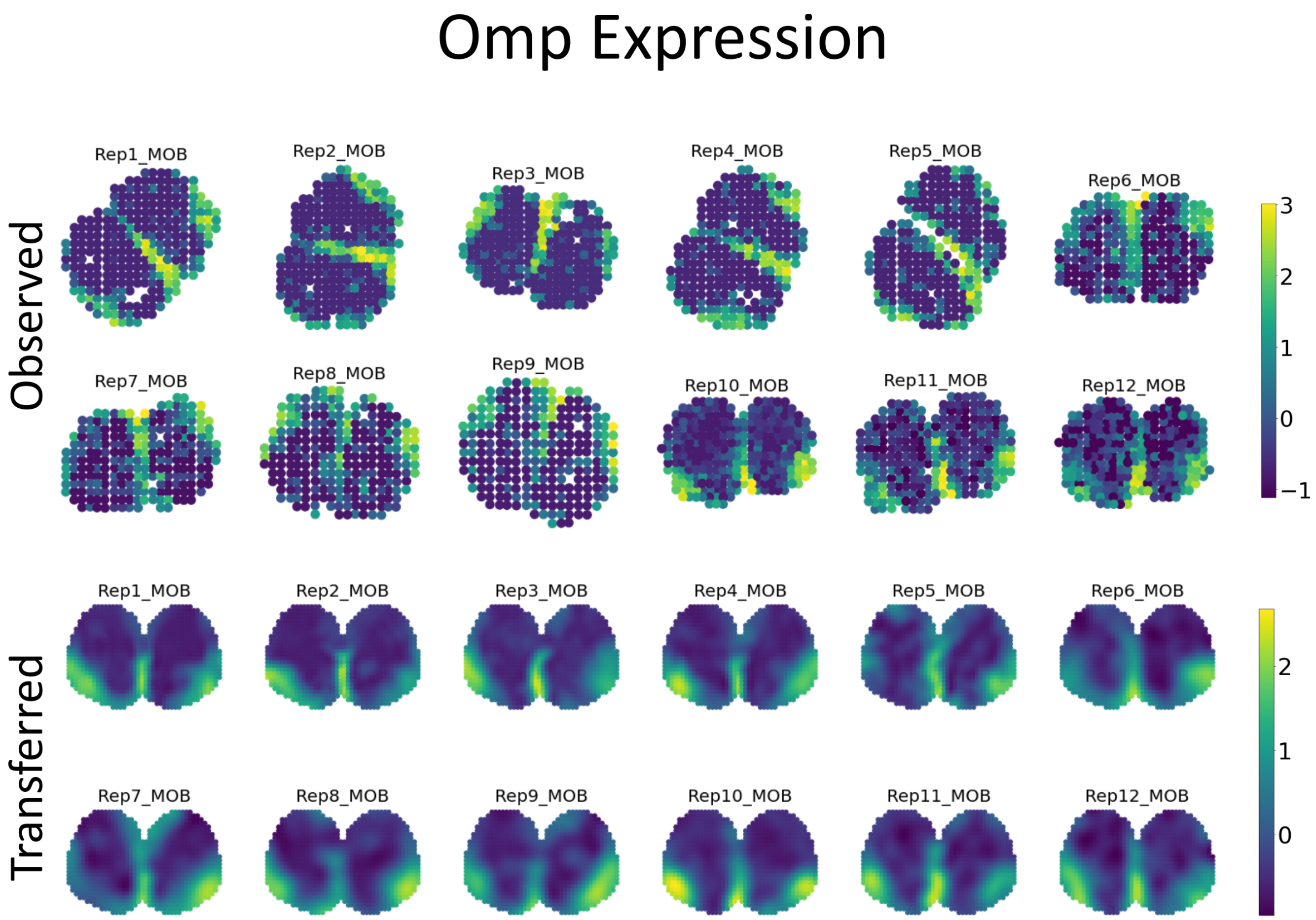


**Supplementary Figure 4:** *Top : Normalized observed spatial expression profiles of Omp in the twelve mouse olfactory bolb (MOB) sections. Bottom : results from transferring the observed spatial gene expression to a shared reference.*

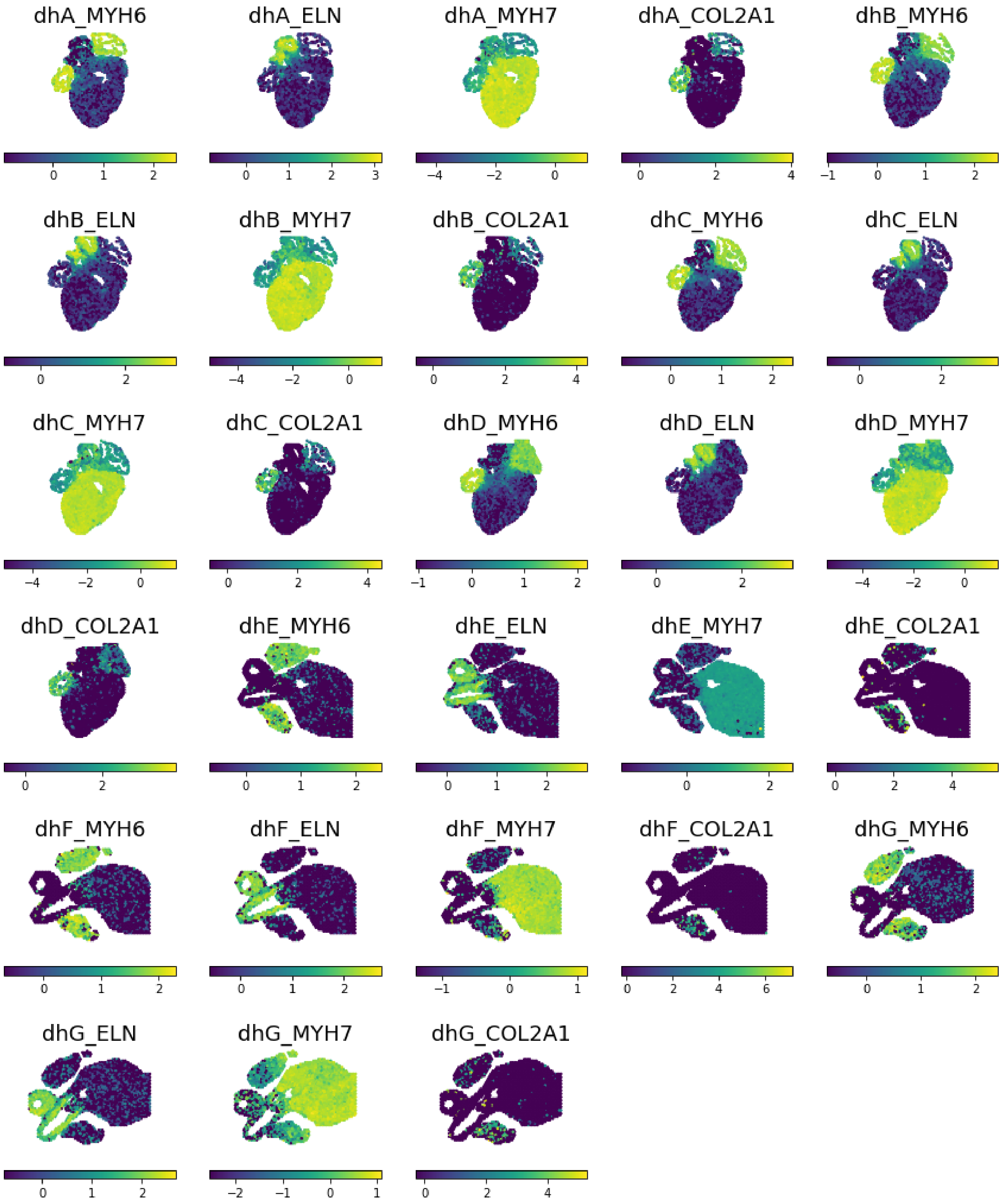


**Supplementary Figure 5:** *Normalized observed spatial expression profiles for all combinations of the seven human developmental heart sections (dhA-D) and the four genes (COL2A1, ELN, MYH6, MYH7).*

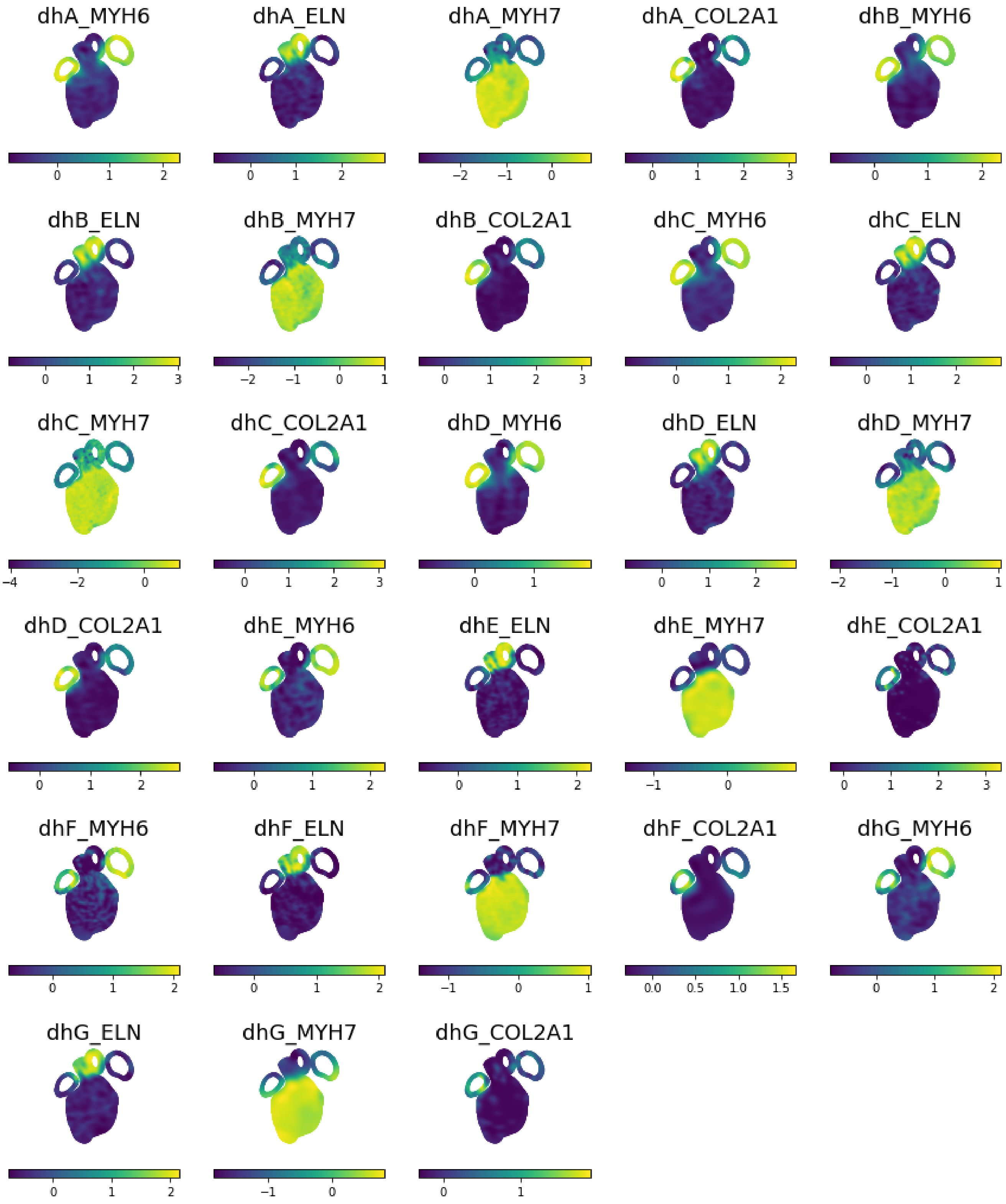


**Supplementary Figure 6:** *Results from transferring the spatial expression profiles of the four different genes COL2A1, ELN, MYH6, MYH7) in the seven human developmental heart sections (dhA-D) to a shared reference.*

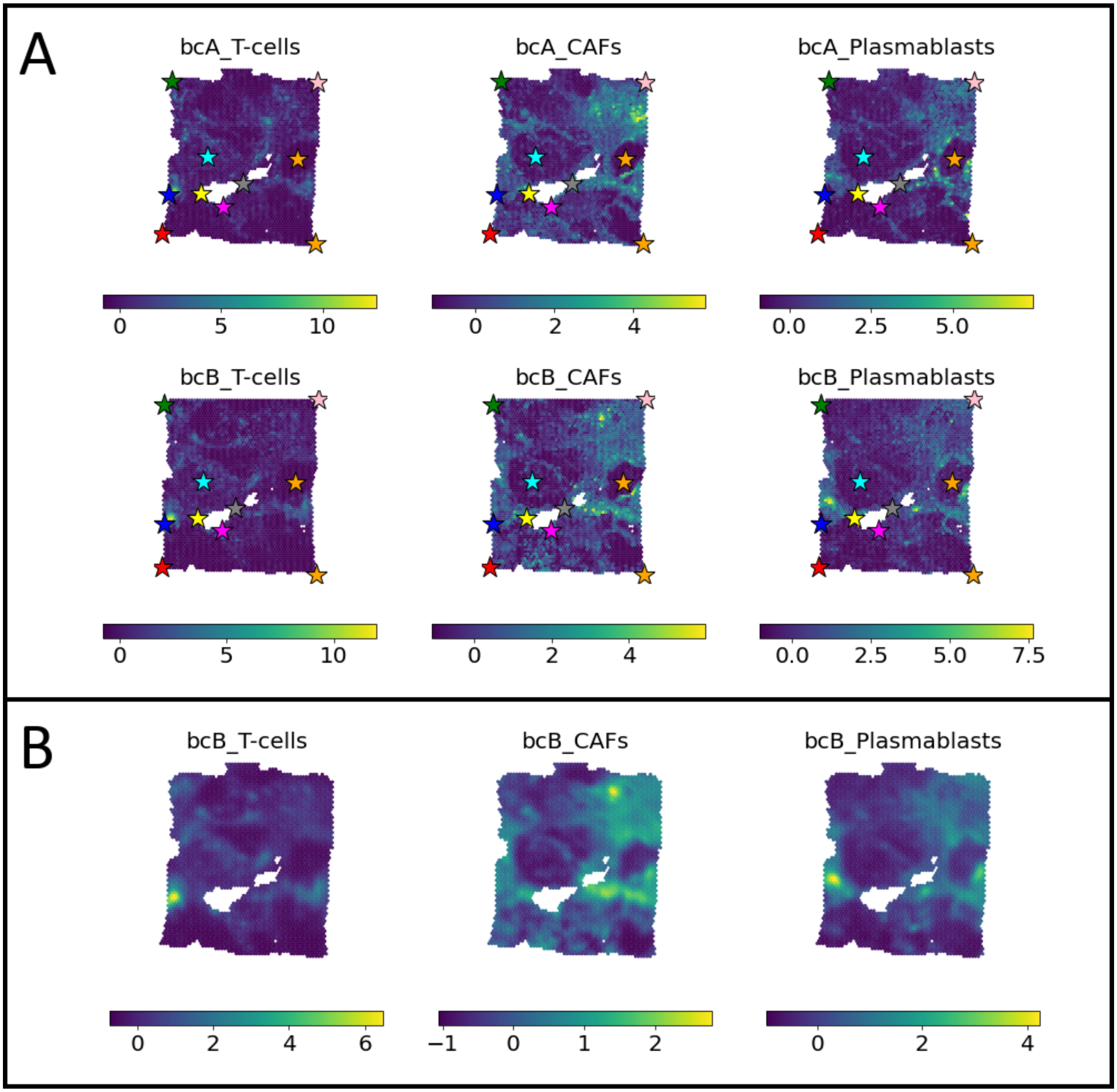


**Supplementary Figure 7:** *We used the tool stereoscope to integrate spatial transcriptomics data from two tissue sections (sample bcA and bcB) of HER2-positive breast cancer surveyed with the Visium platform, and a single cell data set containing cells from HER2-positive patients. This allowed us to decompose the spatial data into proportions of cells belonging to respective cell type at each spatial capture location. Both tissue sections were then charted, and finally we transferred the cell type proportions of T-cells, CAFs and Plasmablasts from bcB to bcA (letting the latter act as a reference).* ***A****) Observed proportion values of each cell type in bcA and bcB.* ***B****) The proportions of bcB cast in the domain of bcA. Here, the transferred values highly resembles the original bcA proportions, only being slightly more smooth in their distribution. To be noted is that bcA and bcB represent two distinct sections of the same tissue, meaning some discrepancies are expected. See Methods for more details regarding the analysis.*

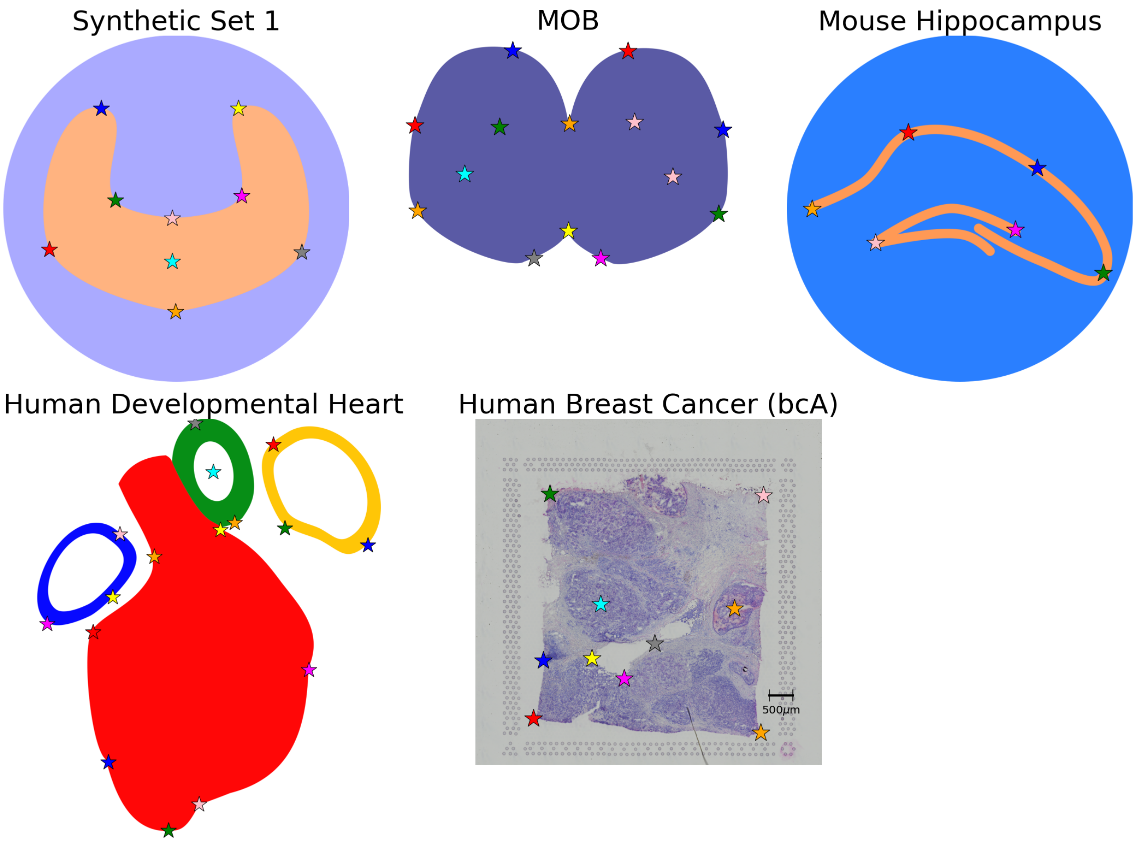


**Supplementary Figure 8:** *Reference templates for the synthetic 1, MOB, hippocampal region, human developmental heart, and human breast cancer data sets. Landmarks are indicated with colored markers.*

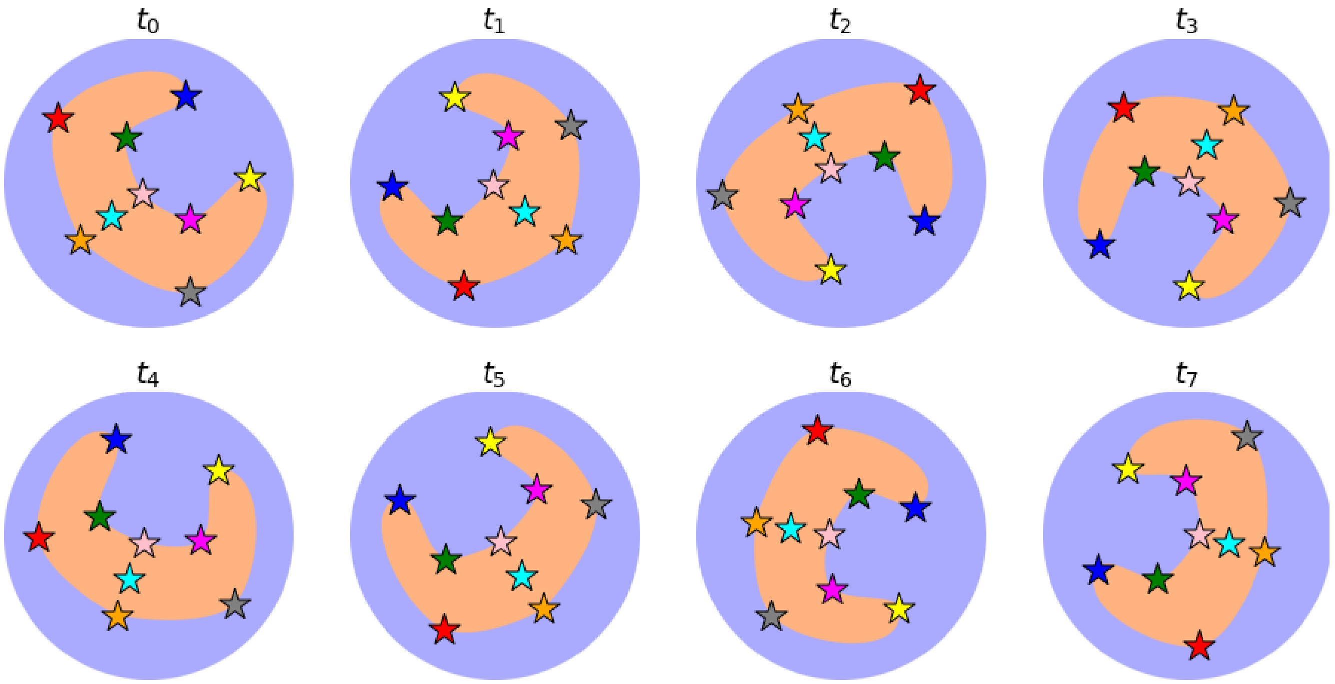


**Supplementary Figure 9:** *Templates for synthetic data set 1 and the related landmarks. Each image was used as a template to generate the "observed" data at the associated time point during the assembly of the synthetic data set, see Methods for more information. Landmarks are indicated with colored markers.*

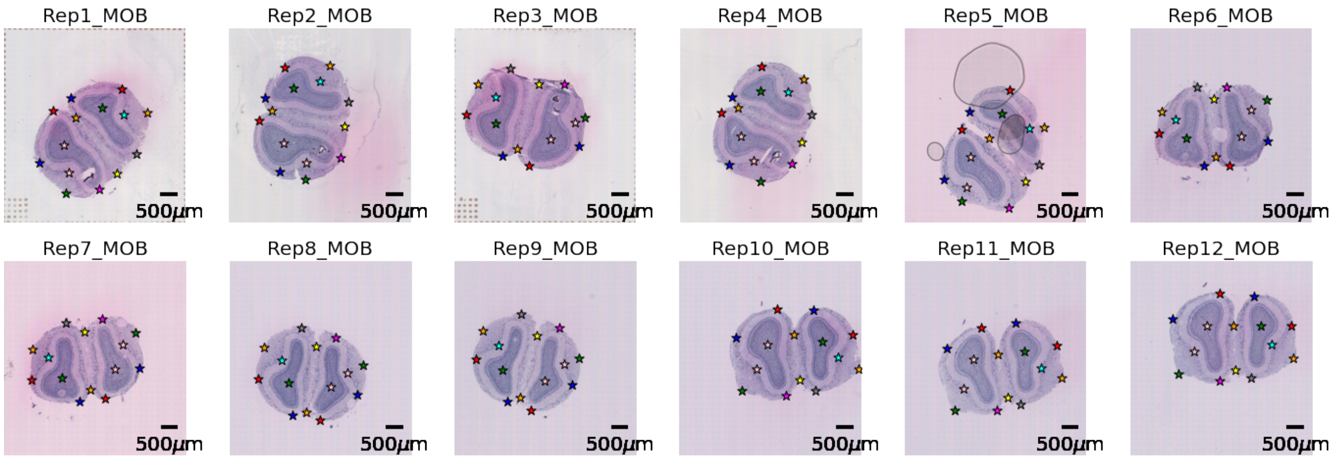


**Supplementary Figure 10:** *The charted MOB data set. Landmarks are indicated with colored markers. The scalebar indicates 500μm.*

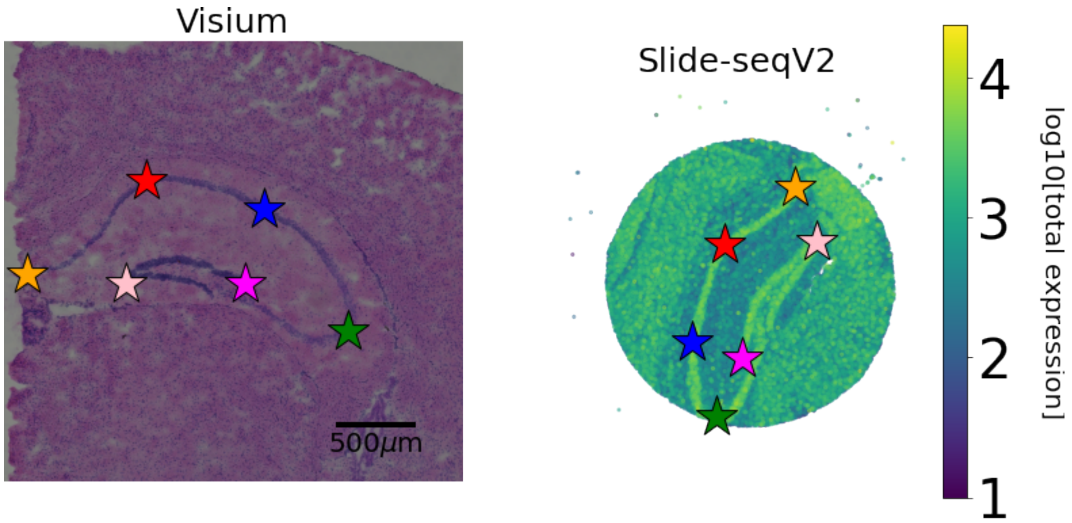


**Supplementary Figure 11:** *The charted Hippocampal region data sets. Landmarks are indicated with colored markers. In the Visium sample the scalebar indicates 500μm. Since the Slide-seqV2 does not provide images of the tissue sample being surveyed we instead represent the tissue by (log10) total UMI count.*

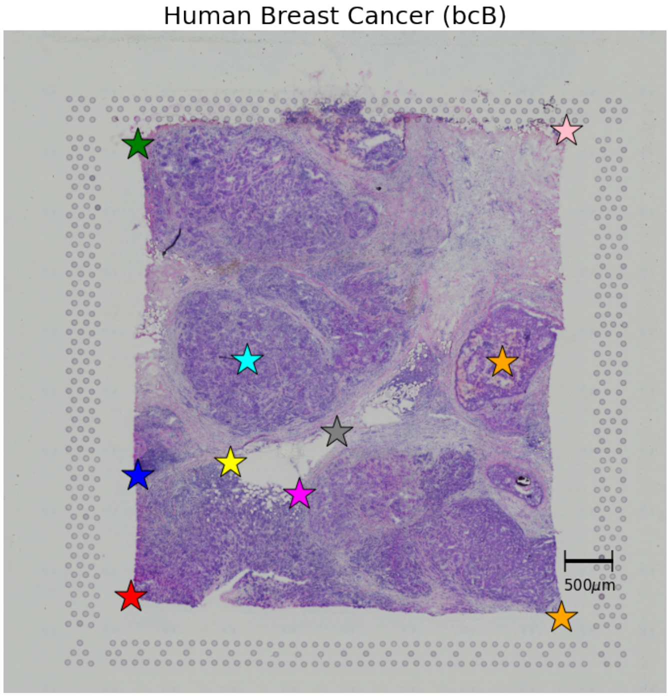


**Supplementary Figure 12:** *The charted bcB sample, landmarks are indicated with colored markers. The scalebar represents* 500*μm.*

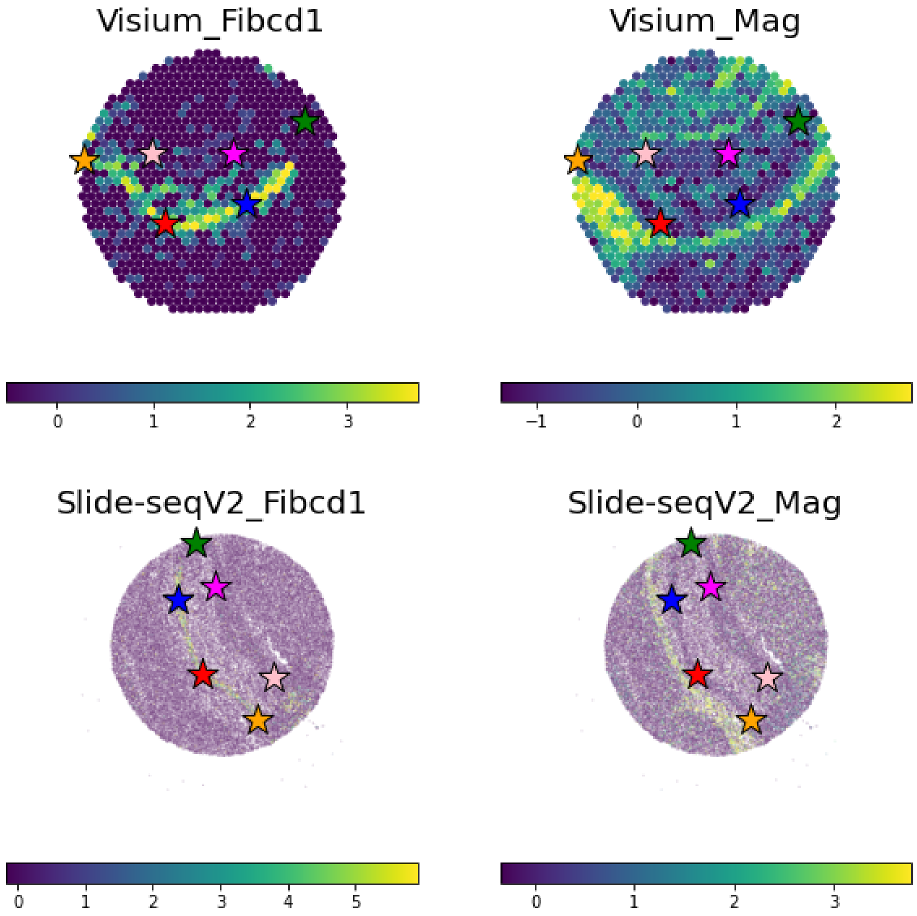


**Supplementary Figure 13:** *The observed gene expression values for Fibcd1 and Mag in the Visium and Slide-seqV2 samples.*

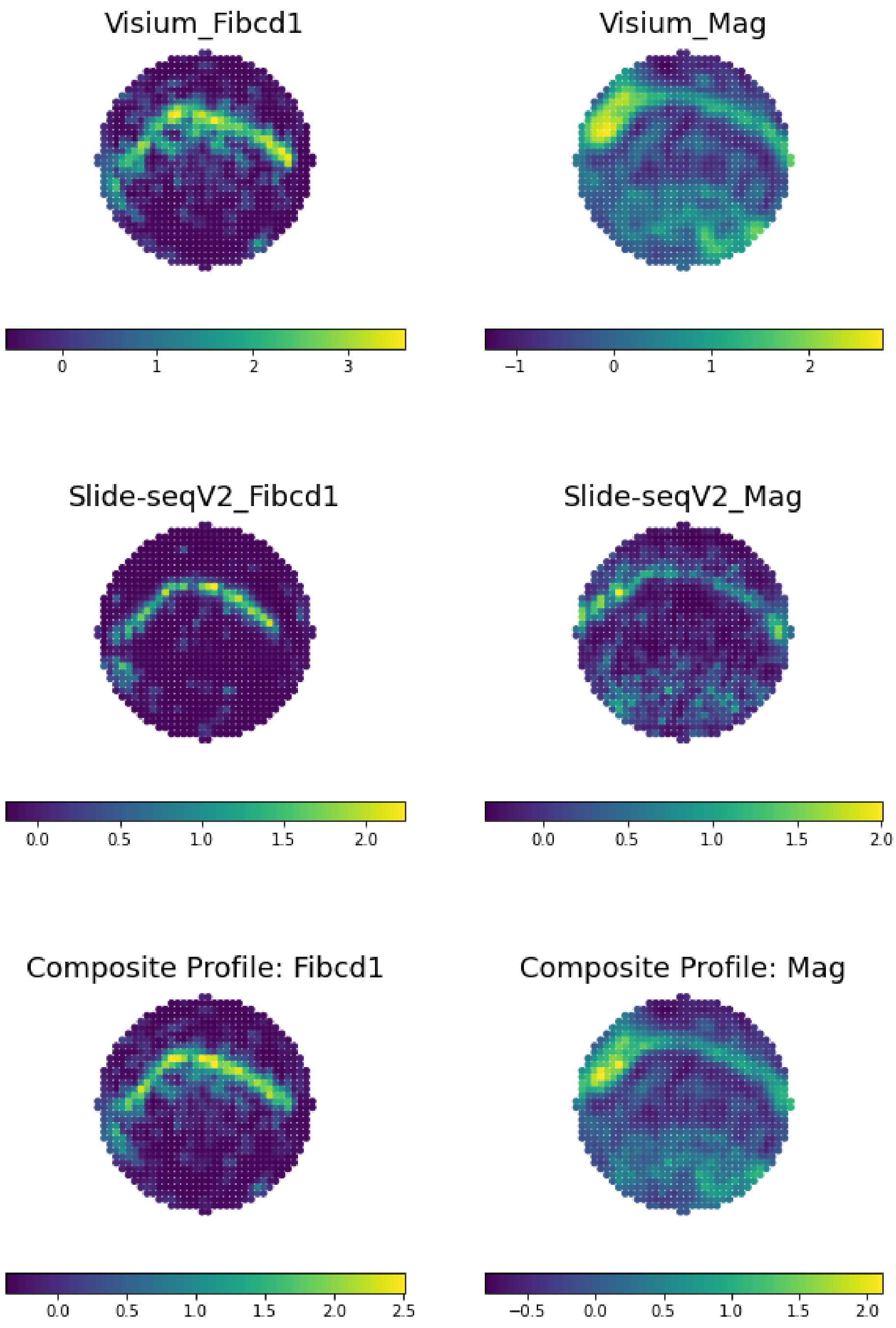


**Supplementary Figure 14:** *The transferred gene expression values for Fibcd1 and Mag in the Visium and Slide-seqV2 samples. The bottom row shows the composite representation (constructed from both samples) for each gene.*

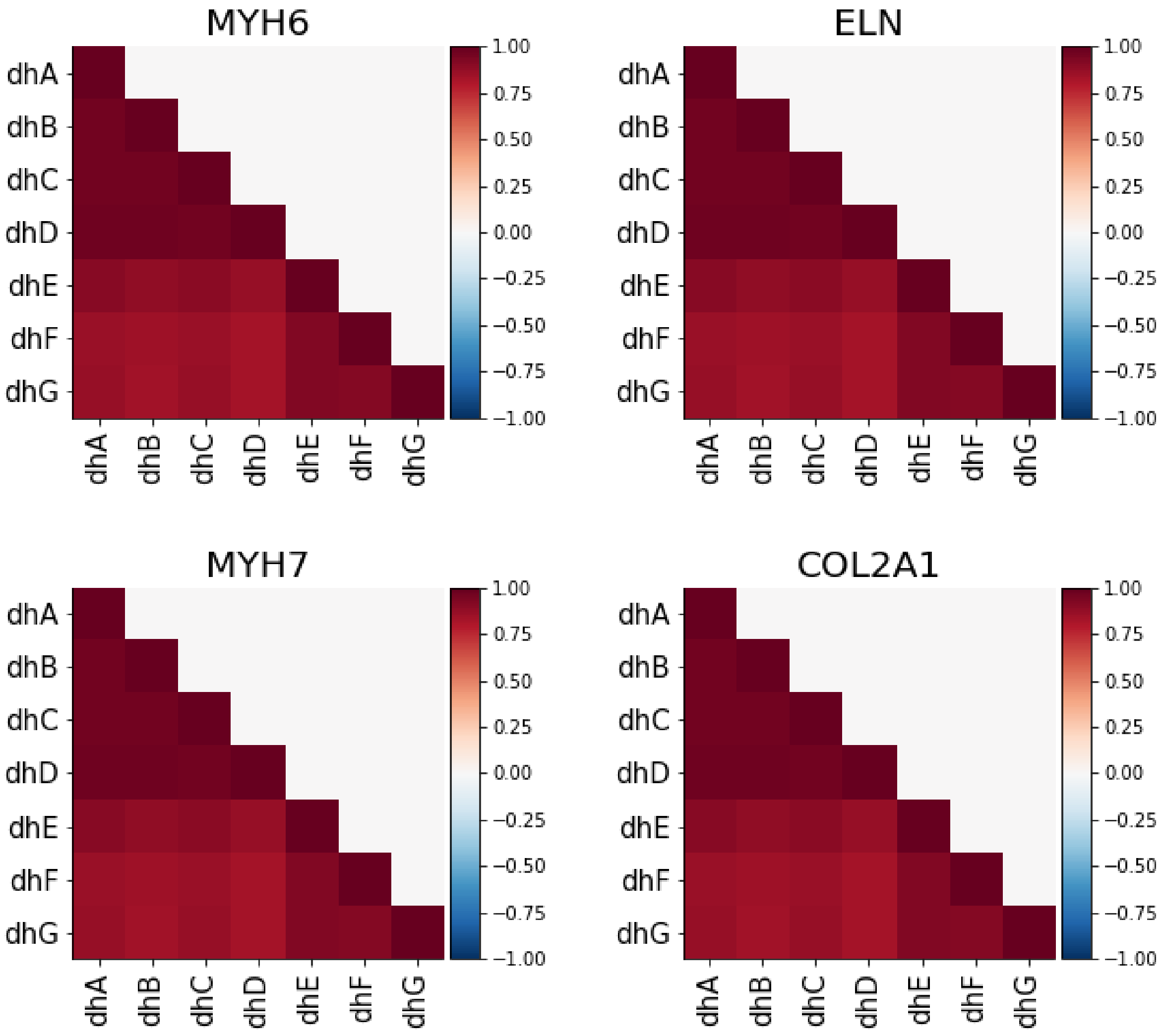


**Supplementary Figure 15:** *Location-wise correlation between all human developmental heart sections (A-F) across the surveyed genes.*

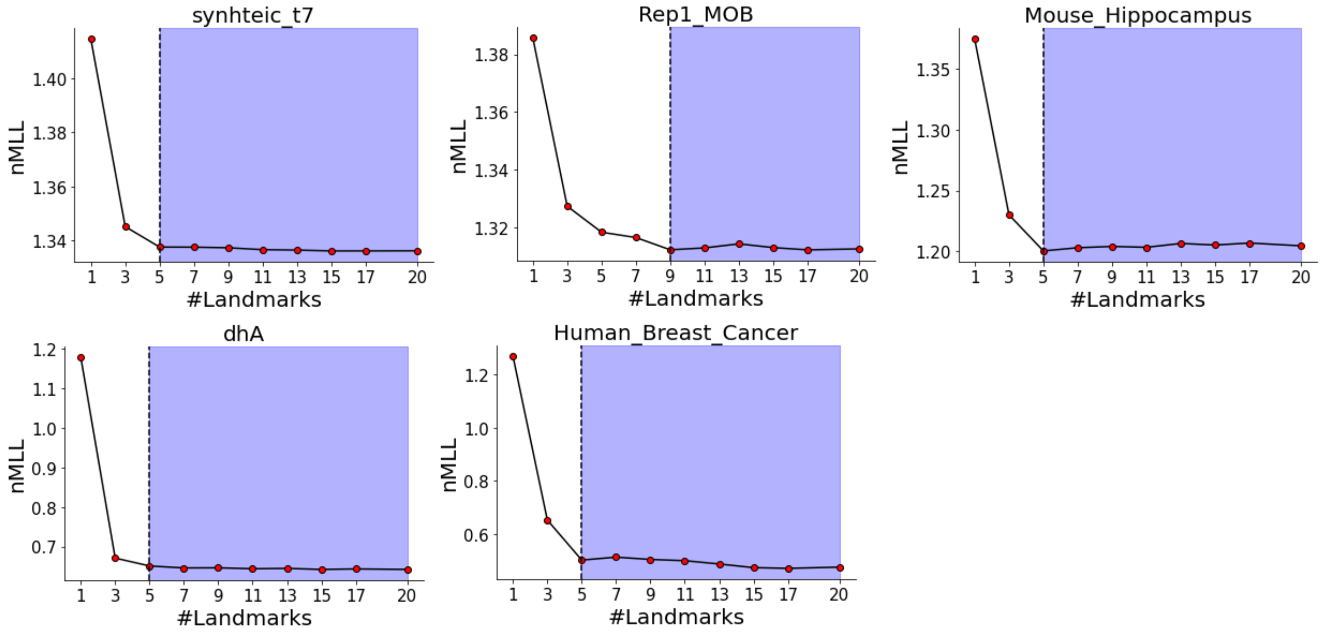


**Supplementary Figure 16:** *Graphs used to determine the lower bound for the number of landmarks in each data set. The dashed black line indicates the lower bound, the blue region shows number of landmarks that are above this lower bound. The negative marginal log likelihood (nMLL) is derived from one representative sample from each data set (synthetic 1: t7, MOB: Rep1, mouse hippocampus: the Visium sample, human developmental heart: dhA, human breast cancer: bcA) and taken as the average over the last 200 epochs ($T = 200$), a process repeated 5 times ($n_{rep} = 5$). In each iteration, the subset of landmarks used to train the model were randomly chosen from a set of 20 randomly positioned landmarks (based on Poisson Disc Sampling). The graphs are smoothed using a Savitzky–Golay filter.*
